# The Telomerase Reverse Transcriptase (TERT) and p53 Regulate Mammalian PNS and CNS Axon Regeneration downstream of c-Myc

**DOI:** 10.1101/547026

**Authors:** Jin-Jin Ma, Ren-Jie Xu, Xin Ju, Wei-Hua Wang, Zong-Ping Luo, Chang-Mei Liu, Lei Yang, Bin Li, Jian-Quan Chen, Bin Meng, Hui-Lin Yang, Feng-Quan Zhou, Saijilafu

## Abstract

Although several genes have been identified to promote axon regeneration in the central nervous system, our understanding of the molecular mechanisms by which mammalian axon regeneration is regulated is still limited and fragmented. Here by using sensory axon and optic nerve regeneration as model systems, we revealed an unexpected role of telomerase reverse transcriptase (TERT) in regulation of axon regeneration. We also provided strong evidence that TERT and p53 acted downstream of c-Myc to control sensory axon regeneration. More importantly, overexpression of p53 in sensory neurons and retinal ganglion cells (RGCs) was sufficient to promote sensory axon and optic never regeneration, respectively. The study revealed a novel c-Myc-TERT-p53 signaling pathway, expanding horizons for novel approaches promoting CNS axon regeneration.

## Introduction

The central reason of failed axon growth in the adult mammalian central nervous system (CNS) is the diminished intrinsic axon regeneration ability in neurons during maturation and aging (Sun and He, 2010). For example, RGC neurons lose their intrinsic axon growth ability during development and maturation (Goldberg et al., 2002). During the past decade, studies focusing on the intrinsic axon growth ability have generated by far the most promising results in the CNS axon regeneration field. For instance, genetic deletion of the PTEN could elevate the intrinsic axon regeneration ability of mature CNS neurons and promote regeneration of optic nerve and corticospinal tract (Liu et al., 2010; Park et al., 2008). Similar molecules regulating the intrinsic axon regeneration ability are KLF4 (Moore et al., 2009), SOCS3 (Smith et al., 2009), SOX11 (Wang et al., 2015), Lin28 (Wang et al., 2018), and c-Myc (Belin et al., 2015) etc. Despite these significant progresses during the past decade, our understanding of the molecular mechanisms by which mammalian CNS axon regeneration is regulated is still fragmented.

In contrast, neurons from the peripheral nervous system (PNS) can regenerate their axons by reactivating the intrinsic axon growth abilities in response to peripheral nerve injury (Chandran et al., 2016; Michaelevski et al., 2010). Such response is mediated by a transcription-dependent process (Saijilafu et al., 2013; Smith and Skene, 1997), in which several transcription factors (TFs) have been identified, such as c-Jun (Raivich et al., 2004; Zhou et al., 2004), Smad1 (Parikh et al., 2011; Saijilafu et al., 2013), ATF3 (Seijffers et al., 2007), and STAT3 (Bareyre et al., 2011; Qiu et al., 2005). Importantly, almost all of these TFs functioned similarly in the CNS to regulate axon regeneration (Bareyre et al., 2011; Fagoe et al., 2015; Finelli et al., 2013; Parikh et al., 2011; Qin et al., 2013). Conversely, most of the aforementioned genes regulating the intrinsic axon regeneration ability of CNS neurons also act to control PNS axon regeneration (Jankowski et al., 2009; Jing et al., 2012; Saijilafu et al., 2013; Wang et al., 2018). Thus, currently it is well-recognized that PNS axon regeneration provides a perfect model system to study the molecular mechanisms underlying mammalian axon regeneration.

The c-Myc proto-oncogene encodes a transcription factor that regulates various physiological activities, such as cell growth, cell proliferation, apoptosis, and cellular metabolism (Dang, 1999). Interestingly, a recent study indicated that overexpression of c-Myc in retinal ganglion cells (RGCs), either before or after optic nerve injury, could promote RGCs survival and axon regeneration (Belin et al., 2015). The study also showed that the expression of c-Myc was significantly increased in sensory neurons following a sciatic nerve lesion. However, the underlying molecular mechanisms by which c-Myc regulates axon regeneration have remain unclear. Previous studies in other non-neuronal systems have identified many downstream target genes regulated by c-Myc. For instance, in some non-neuronal cell lines, c-Myc can transactivate the p53 promoter through its basic-helix-loop-helix (bHLH) recognition motif (Reisman et al., 1993). Moreover, in non-neuronal cells c-Myc has also been shown to be an important transcriptional regulator of telomerase reverse transcriptase (TERT) by directly binding its promoter with other transcription factors (Khattar and Tergaonkar, 2017; Wu et al., 1999). Thus, it is likely that c-Myc might control axon regeneration through these downstream targets. Indeed, during serum induced cell proliferation, c-Myc has been shown to regulate the expression of *ATF3* (Tamura et al., 2014), a well-known regulator of axon regeneration. Additionally, several studies have shown that p53 signaling is necessary for both PNS and optic nerve regeneration (Di Giovanni and Rathore, 2012; Floriddia et al., 2012; Gaub et al., 2011; Gaub et al., 2010; Joshi et al., 2015; Tedeschi et al., 2009). TERT is the catalytic subunit of the enzyme telomerase, which is well known for its role in regulation of telomere extension, cellular aging, and cancer (Maciejowski and de Lange, 2017). In post-mitotic neurons, TERT has been shown to protect neurons from oxidative damages and degenerative changes during aging (Liu et al., 2018; Miwa and Saretzki, 2017; Spilsbury et al., 2015). Inhibition of TERT during iPSC induced neuronal differentiation can produce aged neurons suitable for studying late onset neurodegenerative diseases (Vera et al., 2016). Whether TERT plays any role in regulation of axon growth or regeneration has not been investigated.

In the present study, we demonstrated that c-Myc was both necessary and sufficient for supporting sensory axon growth in vitro and in vivo. Importantly, we provided clear and strong evidence that TERT level was markedly increased in sensory neurons upon peripheral nerve injury. Functionally, inhibition or deletion of TERT in sensory neurons significantly impaired sensory axon regeneration in vitro and in vivo. Moreover, TERT level was regulated by c-Myc in sensory neurons and activation of TERT was able to rescue sensory axon regeneration impaired by down regulation of c-Myc, suggesting that TERT acted downstream of c-Myc to regulate sensory axon regeneration. Lastly, we found that p53 was also up regulated in sensory neurons upon peripheral nerve injury and acted downstream of TERT to regulate sensory axon regeneration. Importantly, overexpression of p53 in sensory neurons and RGCs was sufficient to promote sensory axon regeneration in vivo optic nerve regeneration, respectively. Collectively, our data not only revealed an unexpected function of TERT in regulation of axon regeneration, but also suggested that c-Myc, TERT, p53 signaling might act coordinately to regulate both PNS and CNS axon regeneration.

## Results

### C-Myc is necessary and sufficient for supporting sensory axon regeneration in vitro and in vivo

A recent study demonstrated that peripheral axotomy could upregulate c-Myc expression in adult dorsal root ganglion (DRG) sensory neurons (Belin et al., 2015). However, the functional roles of c-Myc in peripheral axon regeneration remain unknown. Similarly, our immunostaining results showed that c-Myc expression was significantly upregulated in lumbar L4-L5 DRG neurons after sciatic nerve injury (Supplementary Figure S1A, B). Western blot analysis showed that peripheral nerve injury-induced upregulation of c-Myc expression in sensory neurons reached a peak at approximately 3 days post-injury, and then decreased 7 days post-injury (Supplementary Figure S1C, D). The mRNA levels of c-Myc were also increased after peripheral axotomy, indicating enhanced gene transcription (Supplementary Figure S1E). To investigate the functional role of c-Myc in regulation of sensory axon regeneration, we applied the specific c-Myc inhibitor, 10058-F4 (10 μM) to cultured adult DRG neurons for 3 days. The result showed that 10058-F4 significantly impaired the regenerative axon growth (Supplementary Figure S1F, G). Similarly, downregulation of c-Myc expression via a specific small interfering RNA (siRNA) against c-Myc also blocked axon growth of cultured DRG neurons (Supplementary Figure S1H, I), confirming the pharmacological results. We next examined the role of c-Myc in controlling sensory axon regeneration *in vivo* using DRG electroporation technique (Saijilafu et al., 2012; Saijilafu et al., 2014). The specific siRNA against c-Myc, together with the plasmid encoding enhanced green fluorescent protein (EGFP), were co-electroporated into adult mouse L4-L5 DRGs *in vivo.* Two days later, a sciatic nerve crush was performed using fine forceps and the injured site was labeled with 11–0 nylon epineural sutures as previously described Saijilafu et al., 2014. After three more days, the animals were perfused with paraformaldehyde (PFA) and the entire sciatic nerve was dissected out. The resulting EGFP labeled sensory axons were imaged, manually traced, and measured from the injury site to the distal axon tips to quantify axon regeneration by a different person not directly involved in the experiments. The results showed that the siRNA against c-Myc significantly blocked sensory axon regeneration compared to that of control neurons electroporated with a scrambled siRNA and EGFP plasmid (Figure 1A-C).

**Figure 1.**
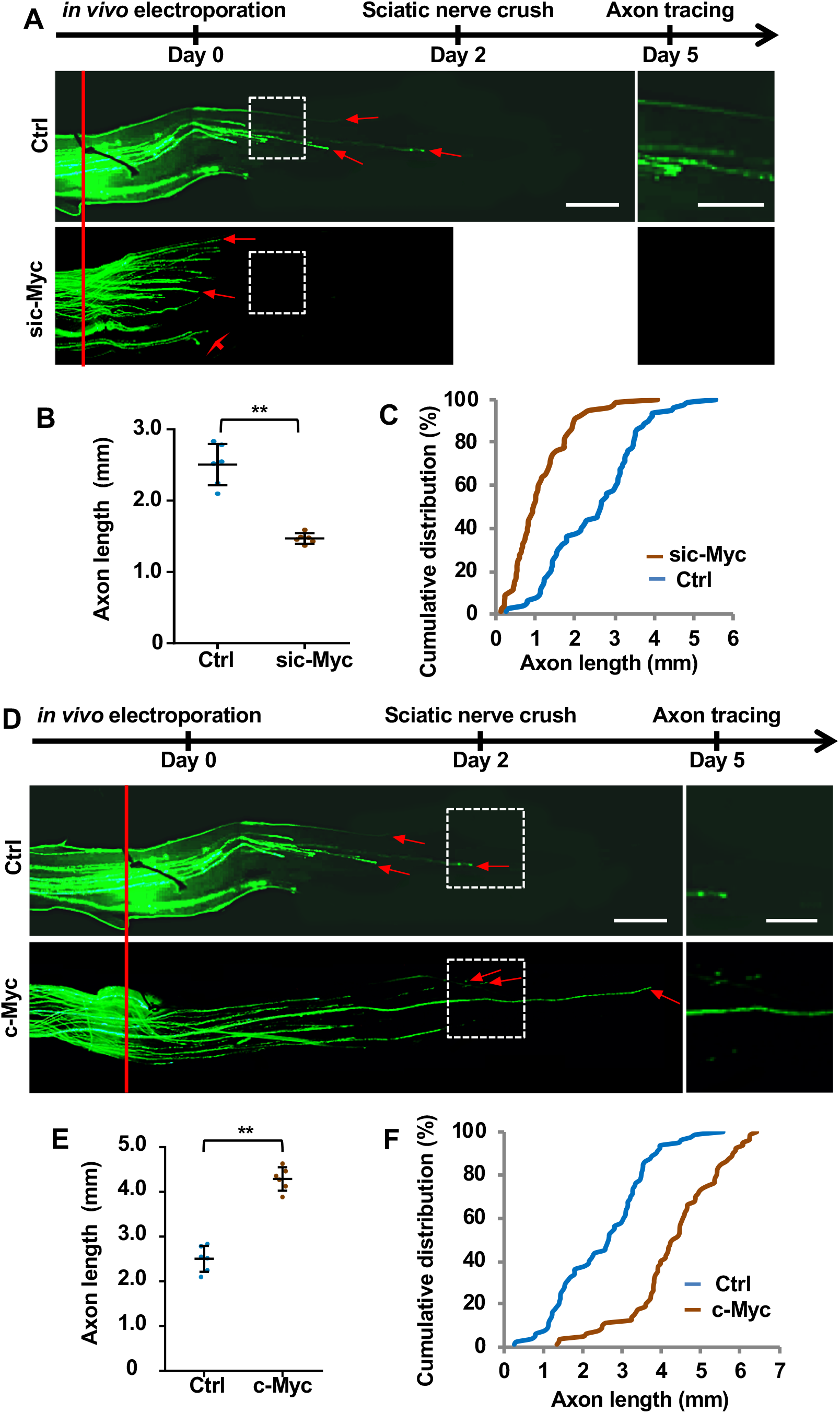
c-Myc is both necessary and sufficient for sensory axon regeneration in vivo. (A) Top: time line of the experiments. Bottom: representative images showing that knocking down c-Myc in sensory neurons in vivo significantly impaired sensory axon regeneration. The red line indicates the nerve crush sites, and the red arrows indicate the distal ends of regenerating axons. The images in the white dotted line boxes in the left panel were enlarged and presented in the right panel. Scale bar: 500 μm in the left panel and 250 μm in the right panel. (B) Quantification of (A) showing that knocking down c-Myc significantly impaired sensory axon regeneration in vivo (n=6 mice for each condition, \*\**p* < 0.01). (C) Cumulative distribution curves showing that knocking down c-Myc in sensory neurons significantly inhibited sensory axon regeneration in vivo. (D) Top: time line of the experiments. Bottom: representative images showing that c-Myc overexpression promoted sensory axon regeneration *in vivo* 3 days after sciatic nerve crush injury. The red line indicates the nerve crush sites and the red arrows indicate distal ends of selected regenerating axons. The images in the white dotted line boxes in the left panel were enlarged and presented in the right panel. Scale bar: 500 μm in the left panel and 250 μm in the right panel. (E) Quantification of (D) showing that overexpression of c-Myc significantly promoted sensory axon regeneration in vivo (n=6 mice for each condition, \*\**p* < 0.01). (F) Cumulative distribution curves showing that overexpression of c-Myc in sensory neurons significantly promoted sensory axon regeneration in vivo. Please note that the control group in A-C is the same as that in D-E.

Overexpression of c-Myc has been shown to increase the intrinsic axon growth ability of RGCs and promote optic nerve regeneration (Belin et al., 2015). We thus tested if c-Myc acted similarly in the PNS to promote axon regeneration. When c-Myc was overexpressed in cultured DRG neurons *in vitro,* we found that sensory axon growth was significantly enhanced on supportive substrates laminin (Supplementary Figure S2A, B). Moreover, CSPGs and myelin are two well-known major inhibitory substrates of axon regeneration (Lee and Zheng, 2012; Ohtake and Li, 2015). We thus investigated if overexpression of c-Myc was able to overcome the inhibitory effects of these substrates. Dissociated sensory neurons transfected with c-Myc plasmid were cultured on CSPGs or myelin coated cell culture plates for 3 days. The results showed that either CSPGs or myelin substrates significantly inhibited the axon growth of DRG neurons, and overexpression of c-Myc markedly enhanced their axon growth on either CSPGs (Supplementary Figure S2C, D) or myelin (Supplementary Figure S2E, F) substrates. To determine the in vivo role of c-Myc in regulation of axon regeneration, we transfected L4-L5 DRGs of adult mouse with the c-Myc plasmid via *in vivo* DRG electroporation, and performed sciatic nerve crush injury two days later. After three days, we found that overexpression of c-Myc significantly promoted sensory axon regeneration in vivo (Figure 1D-F). Taken together, our results provided solid evidence that c-Myc was necessary and sufficient to support sensory axon regeneration in vitro and in vivo.

### TERT functions to regulate sensory axon regeneration in vitro and in vivo

To determine the role of TERT in regulation of sensory axon regeneration, we examined the level of TERT in sensory neurons upon peripheral nerve injury. Both immunostaining and western blot analyses showed that TERT expression in DRG neurons was significantly increased after peripheral axotomy (Figure 2A-D). It should be noted that TERT was mostly localized in the cytoplasm of sensory neurons, which is consistent with previous studies of brain neurons (Liu et al., 2018; Spilsbury et al., 2015). One potential explanation is that in post-mitotic neurons TERT is localized in mitochondria to reduce oxidative stress and protect neurons from apoptosis (Liu et al., 2018). Functionally, inhibition of TERT with a specific pharmacological inhibitor BIBR1532 (Lavanya et al., 2018) impaired regenerative axon growth of sensory neurons in a dose dependent manner (Supplementary Figure S3). Importantly, knocking down TERT with a specific siRNA also effectively blocked sensory axon regeneration *in vitro* (Figure 2E-G), confirming the pharmacological results. To examine if TERT functioned to regulate sensory axon regeneration *in vivo,* by using *in vivo* electroporation technique, we transfected L4-L5 DRG neurons with the specific TERT siRNA, and performed sciatic nerve crush injury two days later. Three days after the nerve crush, sensory axon regeneration was analyzed. We found that knocking down TERT significantly blocked sensory axon regeneration *in vivo* (Figure 2H-J), indicating that TERT was required for sensory axon regeneration. These results revealed an unexpected role of telomere regulatory complex in regulating axon regeneration.

**Figure 2.**
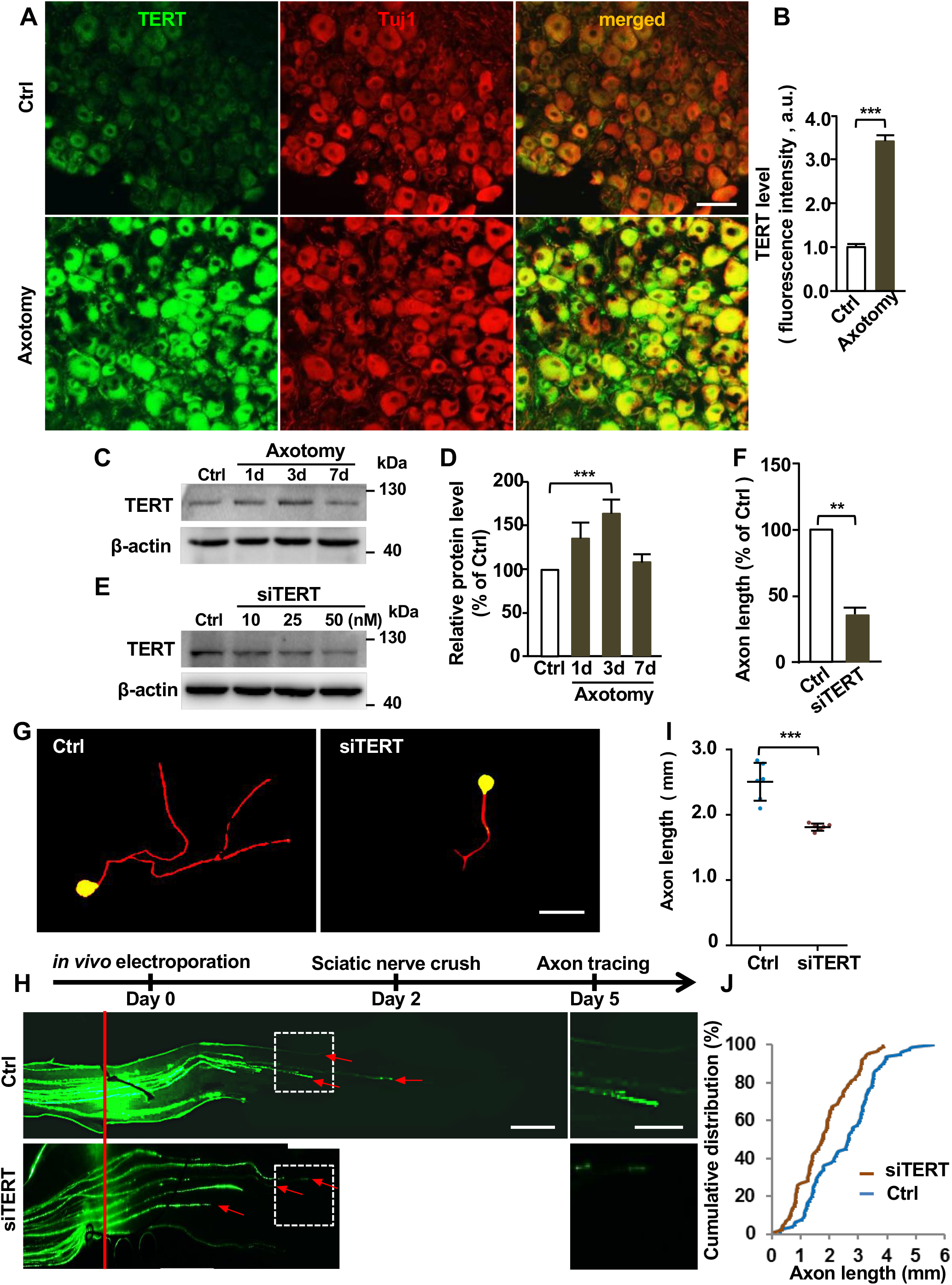
TERT is required for sensory axon regeneration in vitro and in vivo. (A) Representative immunostaining images of sectioned DRGs showing significantly elevated TERT levels in sensory neurons 3 days after sciatic nerve axotomy. Note that most TERT staining in adult sensory neurons was localized in the cytoplasm. Scale bar: 100 μm. (B) Quantification of fluorescence intensity of TERT staining from three independent experiments; \*\*\**p* < 0.001. (C) Representative western blot images showing increased protein levels of TERT in DRG tissues after sciatic nerve axotomy. (D) Quantification of (C) from 3 independent experiments; \*\*\**p* < 0.001. (E) Representative western blot images showing successful knockdown of TERT protein levels in cultured DRG neurons with the siRNA against TERT (siTERT). (F) Quantification of average axon lengths of control neurons or neurons expressing siTERT in cultured sensory neurons from three independent experiments; \*\**p* < 0.01. (G) Representative images showing that knocking down of TERT in cultured sensory neurons with a specific siRNA markedly decreased axon growth after 3 days. Scale bar: 50 μm. (H) Top: time line of the experiment. Bottom: representative images of sensory axon regeneration *in vivo* following TERT knockdown 3 days after sciatic nerve crush injury. The red line indicates the crush sites, and the red arrows indicate the distal ends of selected regenerating axons. The images in the white dotted line boxes in the left panel were enlarged and presented in the right panel. Scale bar: 500 μm in the left panel and 250 μm in the right panel. (I) Quantification of (H) showing that knocking down TERT in sensory neurons significantly inhibited axon regeneration *in vivo* (n=6 mice for each condition, \*\*\**p* < 0.001). (J) Cumulative distribution curves showing that knocking down TERT in sensory neurons significantly reduced sensory axon regeneration in vivo. Please note the control group in H-J is the same as that shown in Figure 1.

To determine if TERT acted downstream of c-Myc in sensory neurons to regulate axon regeneration, we examined the expression level of TERT after knocking down c-Myc with siRNA. The results demonstrated that downregulation of c-Myc resulted in significantly reduced level of TERT in cultured sensory neurons (Figure 3A, B). Conversely, overexpression of c-Myc in sensory neurons markedly enhanced TERT expression (Figure 3C, D). Functionally, when we treated sensory neurons with a specific pharmacological TERT activator CAG (Ip et al., 2014), we found that activation of TERT with CAG alone had no promoting effect on sensory axon growth cultured for 3 days, likely due to upregulation of endogenous TERT. However, CAG (25 μM) treatment significantly rescued regenerative axon growth inhibited by c-Myc knockdown (Figure 3E, F), suggesting that TERT act downstream of c-Myc to regulate axon regeneration.

**Figure 3.**
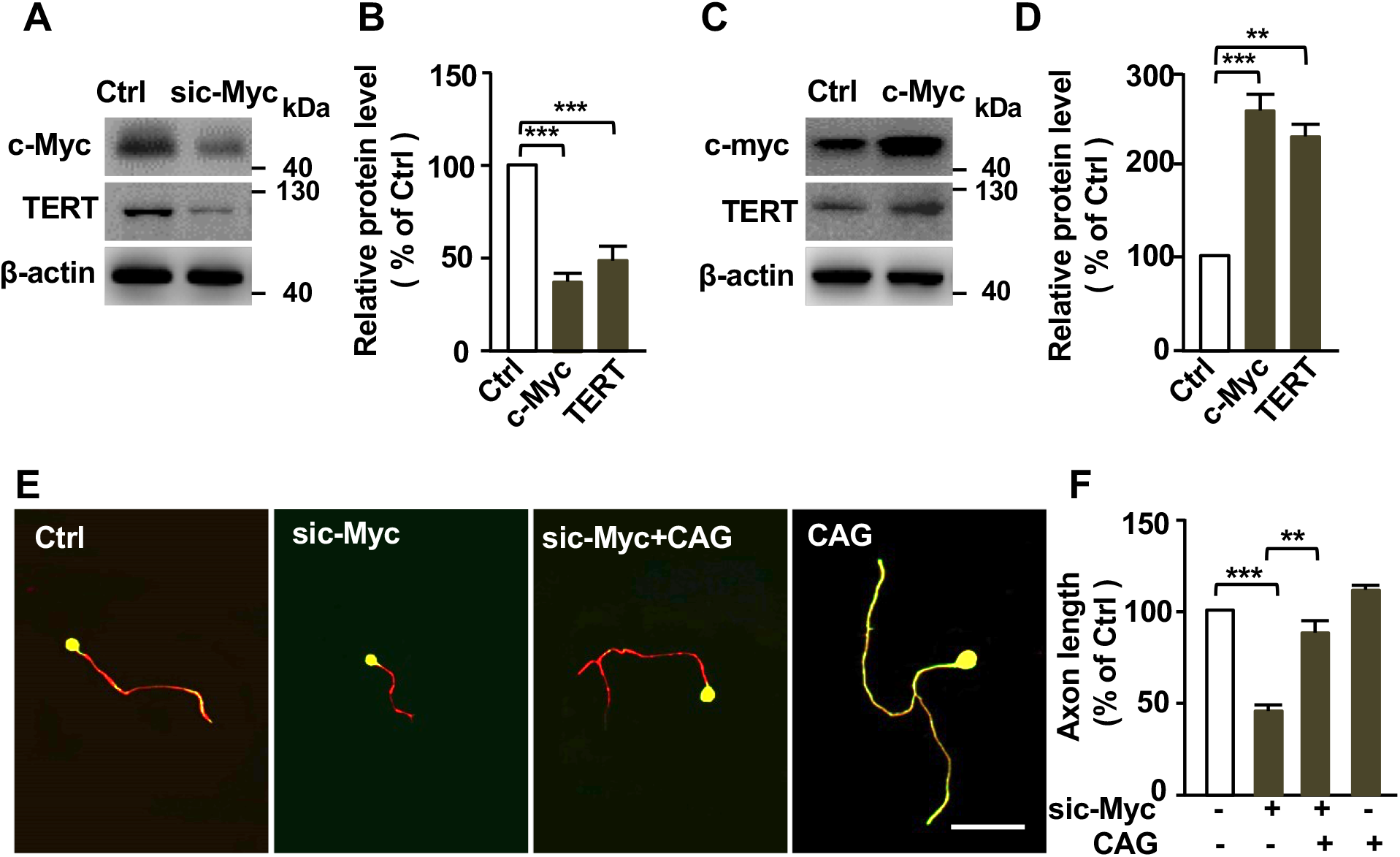
TERT acts downstream of c-Myc in sensory neuron to control axon regeneration. (A) Representative western blot images showing that knocking down c-Myc led to reduced TERT protein levels in sensory neurons. (B) Quantification of (A) showing significantly decreased c-Myc and TERT protein levels in adult sensory neurons after knocking down c-Myc from three independent experiments; \*\*\**p* < 0.001. (C) Representative western blot images showing that overexpression of c-Myc led to increased TERT protein levels in sensory neurons. (D) Quantification of (C) showing significantly increased c-Myc and TERT protein levels of in adult sensory neurons after c-Myc overexpression from three independent experiments; \*\**p* < 0.01, \*\*\**p* < 0.001. (E) Representative images showing that treatment of cultured sensory neurons with the specific TERT activator, CAG (25 μM), for 3 days, successfully rescued sensory axon growth inhibited by c-Myc knockdown. Scale bar: 50 μm. (F) Quantification of average axon lengths of (E) in different conditions from three independent experiments; \*\**p* < 0.01, \*\*\**p* < 0.001.

### p53 is necessary and sufficient for supporting sensory axon regeneration in vitro and in vivo

To investigate if p53 act in the c-Myc-TERT pathway to regulate axon regeneration, we examined the level of p53 in sensory neurons upon peripheral nerve injury. The results showed that p53 was up regulated in sensory neurons following sciatic nerve injury (Figure 4A, B). Functionally, the specific membrane permeable p53 inhibitor, PFTα, markedly inhibited sensory axon growth in a dose dependent manner (Figure 4C, D), indicating that p53 is necessary for regenerative sensory axon growth. These results were consistent with previous studies (Di Giovanni and Rathore, 2012; Floriddia et al., 2012; Gaub et al., 2011; Gaub et al., 2010; Joshi et al., 2015; Tedeschi et al., 2009) that p53 was required for mammalian axon regeneration. Conversely, we showed that overexpression of p53 in cultured sensory neurons significantly promoted axon growth (Figure 4E-G), indicating that p53 alone is sufficient to promote sensory axon regeneration. Together, these results demonstrated that p53 was both necessary and sufficient to support sensory axon regeneration in vitro.

**Figure 4.**
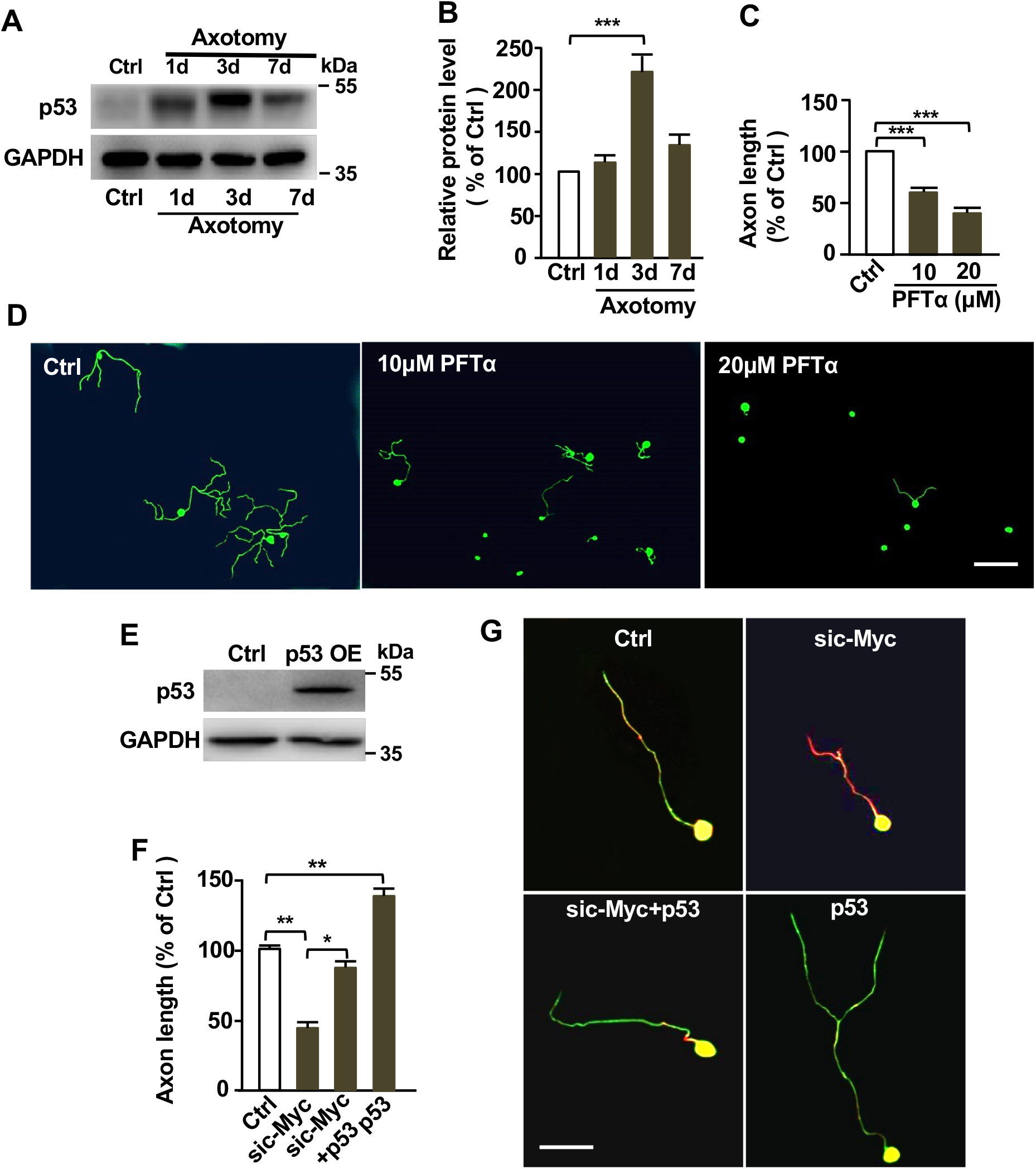
p53 is necessary and sufficient for supporting sensory axon regeneration in vitro. (A) Representative western blot images showing markedly increased p53 protein levels in DRG tissues after sciatic nerve axotomy. (B) Quantification of the western blot images in (A) from three independent experiments; \*\*\**p* < 0.001. (C) Quantification of the average axon lengths in (D) from three independent experiments; \*\*\**p* < 0.001. (D) Representative images showing that inhibition of p53 with PFTα in cultured sensory neurons for 3 days dramatically blocked regenerative axon growth. Scale bar: 200 μm. (E) Representative western blot images showing increased level of p53 after p53 overexpression (OE) in adult sensory neurons. (F) Quantification of the average axon lengths in (G) from three independent experiments; \**p* < 0. 05, **p < 0.01. (G) Representative images showing that overexpression of p53 were sufficient to promote sensory axon regeneration by itself in vitro and restored axon regeneration inhibited by c-Myc knockdown. Scale bar: 50 μm.

Previous studies in non-neuronal cells have shown that p53 activity is controlled by c-Myc (Reisman et al., 1993) or TERT (Zhou et al., 2017). Therefore, we investigated if p53 acted downstream of c-Myc and TERT in sensory neurons to control axon regeneration. Indeed, the specific TERT inhibitor BIBR1532, which has been shown to reduce the protein level of TERT (Lavanya et al., 2018), also led to reduced p53 level in sensory neurons (Supplementary Figure 4A, B), indicating that p53 is regulated by the cMyc-TERT signaling in sensory neurons. Functionally, the p53 activator Tenovin-6 was able to enhance sensory axon growth inhibited by knocking down c-Myc or TERT via siRNA (Supplementary Figure 4C, D), suggesting that p53 act downstream of c-Myc and TERT to regulate axon regeneration. Furthermore, overexpression of p53 significantly enhanced the decreased sensory axon growth caused by c-Myc knockdown in cultured sensory neurons (Figure 4E-G), confirming the pharmacological results.

Lastly, we examined the roles of p53 in regulation of sensory axon regeneration *in vivo* via the electroporation approach. The results showed that overexpression of p53 alone was able to enhance sensory axon regeneration *in vivo* (Figure 5A-C). Moreover, overexpression of p53 significantly restored sensory axon regeneration impaired by c-Myc knockdown. Taken together, our results demonstrated clearly that p53 functioned to regulate sensory axon regeneration *in vitro* and *in vivo,* likely downstream of c-Myc and TERT signals.

**Figure 5.**
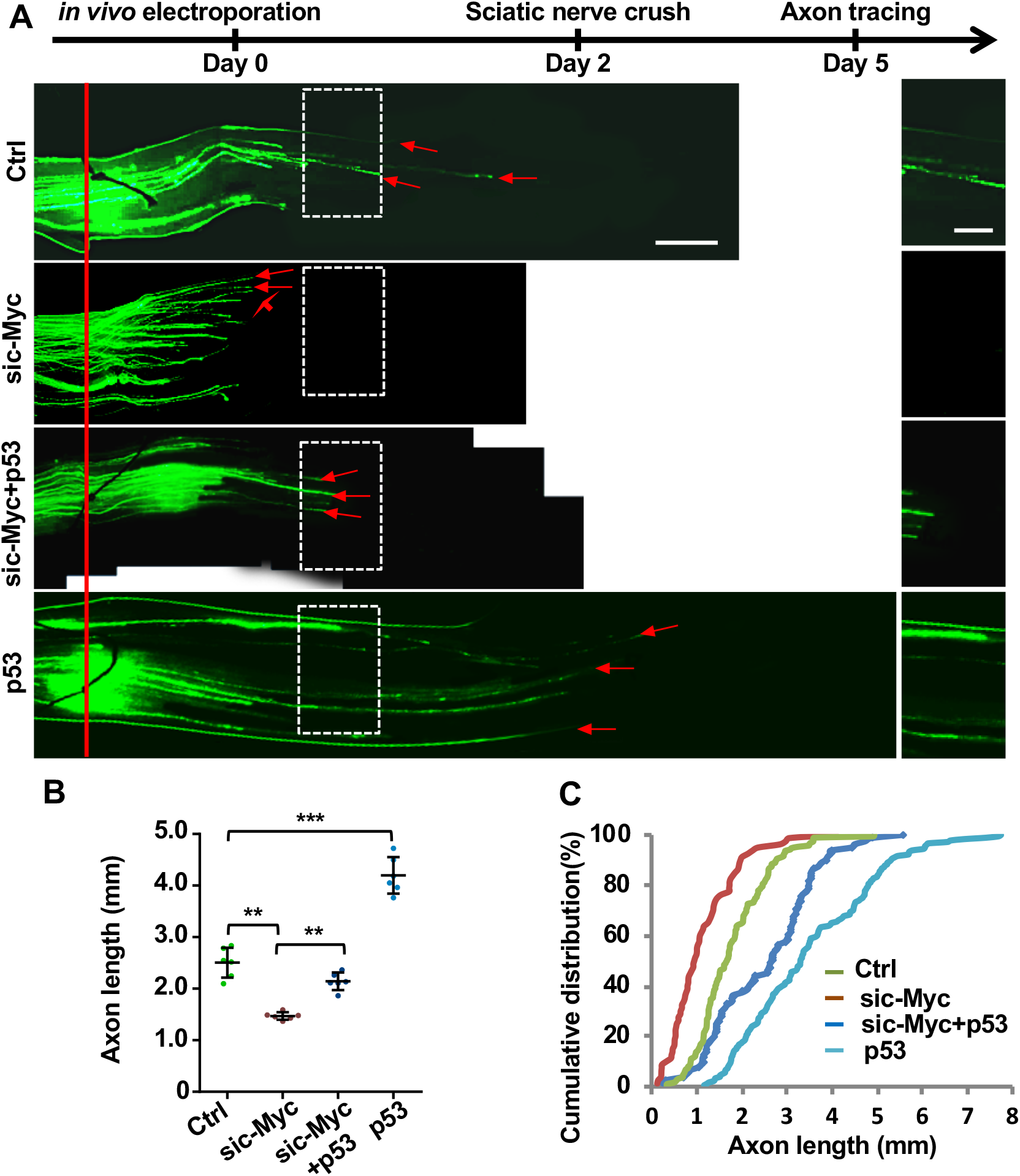
Overexpression of p53 in sensory neurons is sufficient to promote sensory axon regeneration and restore sensory axon regeneration impaired by c-Myc knockdown in vivo. (A) Top: time line of the experiment. Bottom: representative images of sensory axon regeneration in vivo after c-Myc knockdown, overexpression of p53, as well as combined c-Myc knockdown and p53 overexpression. The red line indicates the nerve injury sites and the red arrows indicate the distal ends of selected regenerating axons. The images in the white dotted line boxes in the left panel were enlarged and presented in the right panel. Scale bar: 500 μm in the left panel and 250 μm in the right panel. (B) Quantification of the results in (A) showing that overexpression of p53 alone can promote sensory axon regeneration in vivo and p53 overexpression significantly restores sensory axon regeneration impaired by c-Myc knockdown (n=6 mice for each condition, \*\**p* < 0.01, \*\*\**p* < 0.001). (C) Cumulative distribution curves showing the same results as those in (B). Please note the control and sic-Myc groups in this figure are the same as those shown in Figure 1.

### P53 regulates CNS axon growth and enhances optic nerve regeneration

Our results above showed clearly that p53 was able to enhance mature sensory axon regeneration *in vitro* and *in vivo.* A previous study has shown that p53 is highly expressed in mammalian hippocampal neurons (Qin et al., 2009). Thus, we thought that p53 overexpression might enhance hippocampal axon growth as well. Therefore, we explored the functional roles of p53 in regulating axon growth of embryonic hippocampal or cortical neurons. Embryonic day 18 (E18) hippocampal neurons or E15 cortical neurons were treated with either the p53 inhibitor PFTα or activator Tenovin-6. The results showed that inhibiting p53 with PFTα treatment markedly reduced axonal growth of cortical neurons (Figure 6A, E), whereas activating p53 with Tenovin-6 significantly enhanced cortical axon growth (Figure 6B, F). Similar results were also observed with hippocampal neuron axon growth assay (Figure 6C, D, G, H). These results demonstrated clearly that p53 was also an important regulator of axon growth from embryonic CNS neurons.

**Figure 6.**
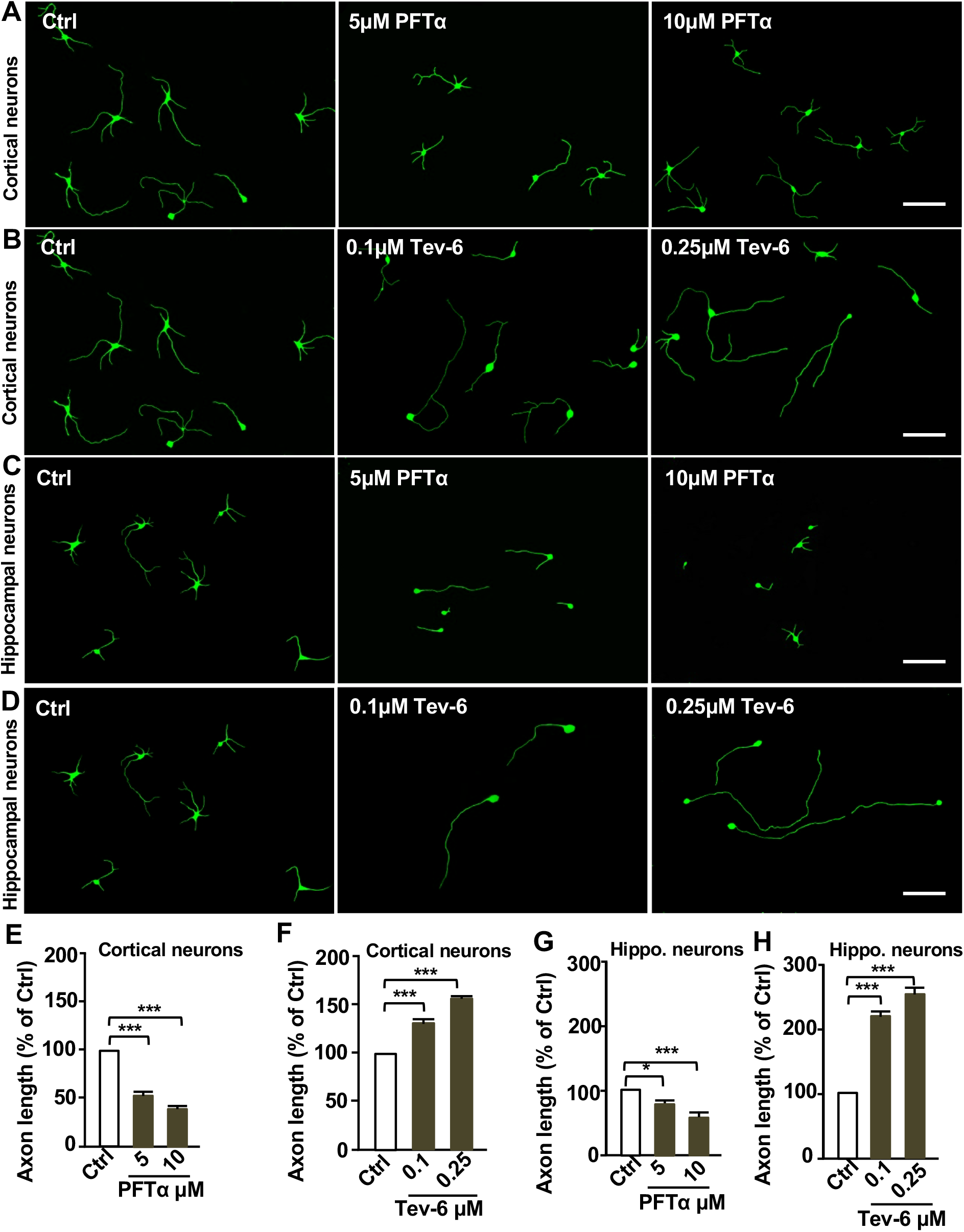
p53 regulates axon growth of developing cortical and hippocampal neurons. (A) Representative images showing that inhibition of p53 activity with the pharmacological inhibitor, PFTα, blocked cortical neuron axon growth. Scale bar: 200 μm. (B) Representative images showing that activation of p53 activity with the small molecule, Tenovin-6, promoted cortical neuronal axon growth. Scale bar: 200 μm. (C) Representative images showing that inhibition of p53 activity with the pharmacological inhibitor, PFTα, blocked hippocampal neuron axon growth. Scale bar: 200 μm. (D) Representative images showing that activation of p53 activity with the small molecule Tenovin-6, promoted hippocampal neuron axon growth. Scale bar, 200 μm. (E) Quantification of the average axon lengths of cortical neurons shown in (A) from three independent experiments; \*\*\**p* < 0.001. (F) Quantification of the average axon lengths of cortical neurons shown in (B) from three independent experiments; \*\*\**p* < 0.001. (G) Quantification of the average axon lengths of hippocampal (Hippo.) neurons shown in (C) from three independent experiments; \**P*<0.05, \*\*\**p* < 0.001. (H) Quantification of the average axon lengths of hippocampal (Hippo.) neurons shown in (D) from three independent experiments; \*\*\**p* < 0.001.

Next, we investigated if p53 overexpression could enhance optic nerve regeneration *in vivo.* Adeno-associated virus 2 (AAV2) viral vector encoding p53 was injected into the vitreous body between the lens and the retina of the eye in a mouse model. The AAV2-GFP viral vectors were used as the control. After two weeks of AAV2-p53 infection, we found that the expression levels of p53 were significantly enhanced in mouse retina tissue (Supplementary Figure S5A) and RGCs (Supplementary Figure S5B, C). To evaluate the functional role of p53 in optic nerve regeneration, optic nerve crush injury was performed two weeks after AAV2-p53 injection. After twelve more days, the fluorescently Alexa-594 conjugated cholera toxin β-subunit (CTB) was injected into the vitreous body to label regenerating axons. The results showed that the number of axons crossing the lesion site in the AAV2-GFP mice two weeks after the optic nerve crush were minimum (Figure 7A, B). In contrast, p53 overexpression induced strong CTB fluorescent labeled regenerating axons (Figure 7A, B). Because p53 is a well-known regulator of cell apoptosis, we assessed the cell viability of RGCs in whole-mount retina immunostained with the neuron-specific tubulin marker, Tuj1. By quantifying the number of cells positive for Tuj1, we found compared to the AAV2-GFP control group, the survival rates of RGCs were no different in the p53 overexpression experimental group, two weeks after the optic nerve crush (Figure 7C, D). These results demonstrated that overexpression of p53 alone was sufficient to promote optic nerve regeneration without affecting RGCs survival.

**Figure 7.**
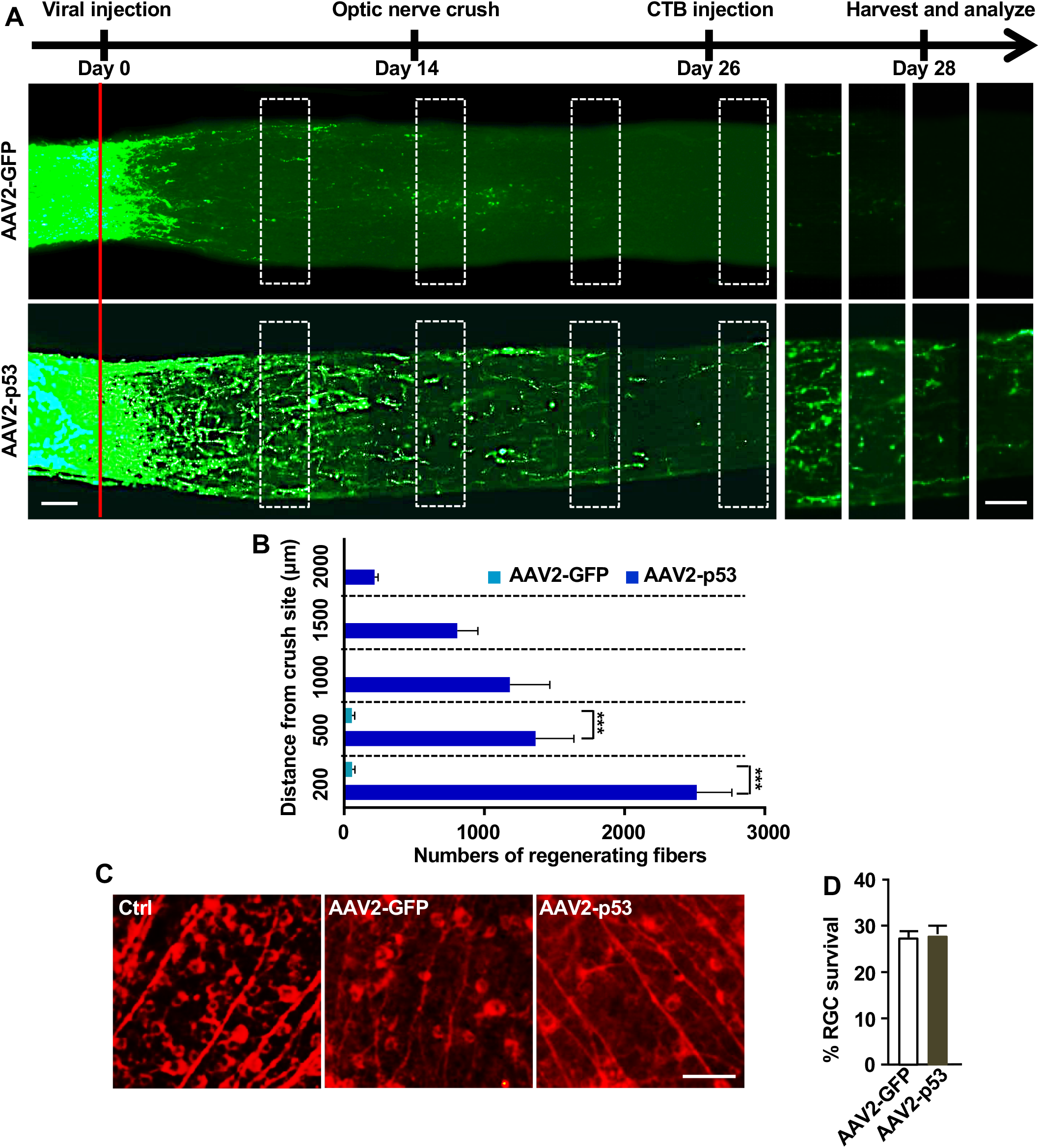
Overexpression of p53 enhances optic nerve regeneration. (A) Top: timeline of the experiment. Bottom: representative images showing that p53 overexpression in RGCs induced drastic optic nerve regeneration 2 weeks after the optic nerve crush. The right 8 columns show enlarged images of nerves at places marked by dashed white boxes on the left. The red line indicates the crush sites. Scale bar: 50 μm for left, 50 μm for right. (B) Quantification of regenerating optic nerve axons at different distances from the nerve crush site (n = 6 mice for each condition, \*\*\**p*<0.001). (C) Representative images of whole mount retina stained with the neuron specific βIII tubulin antibody Tuj-1. Scale bar: 100 μm. (D) Quantification of Tuj-1 positive cells in (C) showing that overexpression of p53 did not affect RGC survival 2 weeks after optic nerve crush.

## Discussion

Identification of novel genes and pathways regulating axon regeneration would not only provide potential new target molecules for treating CNS injuries, but also help us better understand the molecular mechanisms by which mammalian axon regeneration is regulated. Here we demonstrated clearly that c-Myc, previously identified to enhance optic nerve regeneration, was both necessary and sufficient for sensory axon regeneration in vitro and in vivo. Such finding provided a great opportunity and model system to explore novel signaling pathways related to the c-Myc signaling that can be manipulated to regulate mammalian axon regeneration. In this study, we revealed an unexpected role of TERT, a key component of the telomere regulatory complex, in regulating mammalian axon regeneration downstream of c-Myc. Moreover, we provided evidence that p53 likely acted downstream of c-Myc and TERT to regulate sensory axon regeneration. More importantly, we showed that overexpression of p53 alone was sufficient to promote sensory axon regeneration in vivo and optic nerve regeneration.

Besides its well-known function to maintain telomere length during cell division, TERT has been shown to be involved in many other biological functions independent of its telomere regulatory function, such as the regulation of cell metabolism, growth factor secretion, mitochondria function, energy balance, and apoptosis, among others (Bagheri et al., 2006; Chung et al., 2005; Lin et al., 2007; Liu et al., 2012; Massard et al., 2006; Passos et al., 2007; Smith et al., 2003). In post-mitotic neurons, previous studies (Liu et al., 2018; Miwa and Saretzki, 2017; Spilsbury et al., 2015) have shown that TERT is mainly localized in the cytoplasm and function to protect neurons from oxidative or degenerative stresses in aged neurons. It is likely that TERT acts in neurons to regulate mitochondria function by reducing reactive oxidative species (ROS) (Liu et al., 2018; Miwa and Saretzki, 2017; Spilsbury et al., 2015). In this study, we observed that TERT was also mainly localized in the cytoplasm of sensory neurons, similarly to that observed in previous studies, and the expression of TERT was significantly increased in sensory neurons after peripheral injury. Functionally, TERT was required for the sensory axon regeneration in vitro and in vivo. TERT has been shown to be regulated by multiple positive and negative regulators, such as c-Myc, Sp1, p53, Wilms tumor 1 (WT1), E2F, (Cukusic et al., 2008), and the nuclear factor kappa B (NF-ĸB) (Lingner et al., 1997; Nakamura et al., 1997; Yin et al., 2000). Here we showed that in adult sensory neurons down regulation of c-Myc significantly reduced TERT expression, whereas the upregulation of c-Myc significantly increased TERT expression. Functionally, pharmacological activation of TERT significantly rescued the axon regeneration impaired by c-Myc knockdown in adult sensory neurons. Thus, our data strongly suggest that TERT act as a downstream target of c-Myc during sensory axon regeneration.

The p53 protein is well-known for its tumor suppressor role, functioning primarily to promote cell cycle arrest, DNA repair, and apoptosis (Levine and Oren, 2009). In post-mitotic neurons, p53 has been shown to be necessary for proper axon guidance, growth, and regeneration (Arakawa, 2005;Di Giovanni and Rathore, 2012; Floriddia et al., 2012; Gaub et al., 2011; Gaub et al., 2010; Joshi et al., 2015; Tedeschi et al., 2009). However, if p53 is sufficient to promote axon growth and regeneration is unclear. In this study, we showed that overexpression of p53 in adult sensory neurons was sufficient to promote sensory axon regeneration in vitro and in vivo. In addition, the results showed that the protein level p53 was regulated by TERT, suggesting that p53 act downstream of c-Myc and TERT to regulate sensory axon regeneration. Indeed, pharmacological activation of p53 was able to significantly restore sensory axon regeneration *in vitro* impaired by either c-Myc or TERT knockdown. Moreover, overexpression of p53 could significantly restore sensory axon regeneration impaired by c-Myc knockdown both *in vitro* and *in vivo,* providing strong evidence that p53 acted downstream of c-Myc and TERT to regulate axon regeneration. Besides peripheral sensory neurons, we also showed that p53 was an important regulator of axon growth of developing cortical or hippocampal neurons, indicating its similar function in controlling CNS axon growth. Most importantly, we showed clearly that overexpression of p53 in RGCs was sufficient to promote optic nerve regeneration.

How c-Myc, TERT and p53 regulate axon regeneration remains unclear. One potential mechanism is through regulation of cell metabolism. In support, c-Myc is well known to enhance anabolic metabolism of glycolysis via reprogramming mitochondria (Ward and Thompson, 2012). A latest study (Kim et al., 2019) also showed that wild type p53 can act in the mitochondria to suppress oxidative phosphorylation and thus increase glycolysis, similar to that of c-Myc. As aforementioned, TERT has been shown to be localized in mitochondria of neurons and regulates mitochondria function. It would be interesting in the future to explore if other factors or pathways regulating neuronal metabolism could also promote axon regeneration.

## Materials and methods

### Animals and Surgical Procedures

Institute of Cancer Research (ICR) adult female mice, 8 to 12-week-old, (weighing 25 grams [g] to 35 g) were used. All animals were handled according to the guidelines of the Institutional Animal Care and Use Committee of the Soochow University, Suzhou, Jiangsu, China. For surgical procedures, mice were anesthetized with a mixture of ketamine (100 mg/kg) and xylazine (10 mg/kg) via intra-peritoneal (IP) injection. Eye ointment containing atropine sulfate was applied to protect the cornea during surgery, and animals received antibiotics for 24 hours as a post-operative analgesic.

### Reagents and Antibodies

10058-F4, BIBR1532, PFTα, and Tenovin-6 were from Selleck Chemicals (Houston, TX, USA), and CAG was from Sigma-Aldrich (Saint Louis, MO, USA). Antibodies against the neuron-specific class III β-tubulin mouse mAb (Tuj1, 1:1000) were from Sigma-Aldrich. The antibody against c-Myc rabbit mAb (1:1000) was from Genetex (Irvine, CA, USA). Antibodies against TERT rabbit mAb (1:1000) and p53 mouse mAb (1:1000) were from Abcam (Cambridge, UK). The siRNA against TERT and c-Myc were from Gene Pharma (Shanghai GenePharma Co.; Shanghai, China). pEX4-c-Myc and pEX3-p53 plasmids were from Gene Pharma. The AAV2-p53 viral vector was purchased from Cyagen Biosciences (Santa Clara, CA, USA). All fluorescence secondary antibodies were purchased from Molecular Probes, Inc. (Eugene, OR, USA).

### Cell Cultures and Axon Length Quantification

Dissection and culture of adult DRG sensory neurons were performed as described in our previous protocol (Saijilafu et al., 2013). Briefly, DRGs were dissected out from 8-week old adult mice and incubated with collagenase A (1 mg/ml; Roche, Basel, Switzerland) for 90 minutes and then with 1X TrypLE (Life Technologies; Carlsbad, CA, USA) for 20 minutes at 37°C. Then, DRGs were dissociated in culture medium, which was minimum essential media (MEM) supplemented with 5% fetal bovine serum (FBS), antimitotic agents (20 μM 5-fluoro-2-deoxyuridine, 20 μM uridine) and penicillin / streptomycin. The isolated neurons were then centrifuged down and plated onto glass coverslips, which were coated with poly-D-lysine (100 μg/ml, Sigma Aldrich) and laminin (10 μg/ml, Sigma Aldrich). For DRG assays on CSPGs or myelin, culture coverslips were coated with 70 μL CSPGs (5 μg/ml) or purified CNS myelin, which contain a number of axon growth inhibitors. Dissociated DRG neurons were plated onto plastic coverslips in 24-well plates after washing and grown in culture medium (MEM in 5% fetal bovine serum, 20 μM 5-fluoro-2-deoxyuridine, 20 μM uridine, 100 units /ml penicillin and 100 ng/ml streptomycin, Sigma Aldrich) for 3 days at 37°C in a CO2-humidified incubator. For DRG neurons obtained from mice used in axotomy procedures, the culture time for axon growth was 20 hours. Cortical and hippocampal neurons were isolated from embryonic day 15 or 18 (E15, E18) ICR mice embryos and treated with TrypLE for 5 minutes at 37°C. The supernatants were cultured in neurobasal medium supplemented with penicillin / streptomycin, GlutaMAX supplements, and B27 supplements for 3 days.

All images were analyzed with the AxioVision 4.7 software (Carl Zeiss MicroImaging, Inc.; Jena, Germany). In each experiment, at least 100 neurons per condition were selected randomly, and three independent experiments were performed to quantify the average axon length. The longest axon of each neuron was traced manually using the “measure / curve” application.

### RNA Interference

In order to transfect siRNA and / or DNA plasmids into DRG neurons, the dissociated neurons were centrifuged down to remove the supernatant and resuspended in 80 to 100 μl of Amaxa electroporation buffer for mouse neurons (Lonza Cologne GmbH; Cologne, Germany) with mixtures of siRNA (0.2 nmol per transfection) or plasmids and / or the EGFP plasmid (5 μg per transfection). The suspended cells were then transferred to a 2.0-mm cuvette, followed by electroporation with the Amaxa Nucleofector apparatus. After electroporation, cells were immediately mixed with the desired volume of pre-warmed culture medium and plated onto culture dishes coated with a mixture of poly-D-lysine (100 μg/ml) and laminin (10 μg/ml). After the neurons were fully attached to the coverslips (4 hours), the culture medium was changed to remove the remnant electroporation buffer. After 3 days of culture, the cells were fixed with 4% paraformaldehyde (PFA) and then subjected to immunocytochemistry analysis.

### Myelin Extract

Myelin fractions were isolated as described by Norton and colleagues (Norton and Poduslo, 1973). Briefly, three carefully cleaned adult rat brains were homogenized with a Dounce homogenizer in approximately 20 vol. (w/v) of 0.32 M sucrose. We used three brains per 100 ml. One-third of the homogenate was layered over 25 ml of 0.85 M sucrose in each of three tubes, and the tubes were centrifuged at 75,000 g (25,000 rev/min) for 30 minutes. Layers of crude myelin which formed at the interface of the two sucrose solutions were collected with a Pasteur pipette, everything else was discarded. The combined myelin layers were then suspended in water by homogenization and brought to a final volume of 180 ml. This suspension was centrifuged at 75,000 g (25,000 rev/min) for 15 minutes. The supernatant fluid was subsequently discarded. The three crude myelin pellets were again dispersed in a total volume of 180 ml of water and centrifuged at 12,000 g (10,000 rev/min) for 10 minutes. The cloudy supernatant fluid was discarded. The loosely packed pellets were again dispersed in water and centrifuged again. The myelin pellets were then combined and suspended in 100 ml of 0.32 M sucrose. This suspension was layered over 0.85 M sucrose in three tubes and centrifuged exactly as previously described Norton and Poduslo, 1973. The purified myelin was finally removed from the interface with a Pasteur pipette.

### Immunohistochemistry

Mice were deeply anesthetized and perfused with phosphate buffered saline (PBS) and ice-cold 4% PFA. The tissues of lumbar L4-L5 DRG neurons were dissect out and post fixed with 4% PFA at 4°C overnight, followed by dehydration in 10%, 20%, and 30% sucrose solution (w/v) overnight, respectively. DRGs were then cryosectioned into slices, 12 μm in thickness. For immunostaining, the sections of DRG neurons were then washed three times in PBS containing 0.5% TritonX-100, followed by blocking in PBS containing 5% FBS and 0.3% Triton X-100 for 1 hour at room temperature. Sections were incubated with the indicated primary antibodies overnight at 4°C, washed in PBS containing 0.5% TritonX-100, and then incubated with the corresponding secondary antibodies for 1 hour at room temperature. The sections were mounted with mounting medium (Vector Labs, H-1400) after washing in PBS containing 0.3% Triton X-100.

### RNA Isolation and qRT-PCR

The total RNA was extracted from the DRG neurons using the TRIzol reagent (Invitrogen; Carlsbad, CA, USA). The RNA was reverse transcribed into cDNA using Maxima H Minus Reverse Transcriptase according to the manufacturer’s instructions (Thermo Scientific, Waltham, MA, USA). Quantitative PCR (qPCR) was carried out using SYBR-Green Real-Time PCR Master Mix (Toyobo Co.; Osaka, Japan). Standard curves (cycle threshold values versus template concentration) were prepared for each target gene and the endogenous reference (18S) in each sample. The primer sequences were as follows; c-Myc: forward, 5’-ATCACAGCCCTCACTCAC-3’ and reverse, 5’-ACAGATTCC ACAAGGTGC -3’; TERT: forward, 5’-TGGTGGAGGTTGCCAA-3’ and reverse, 5’-CCACTGCATACTGGCGGATAC-3’; p53: forward, 5’-CGACGACATTCGGAT AAG-3’ and reverse, 5’-TTGCCAGATGAGGGACTA-3’. The amplification step involved an initial denaturation at 95°C for 10 seconds (s) followed by 40 cycles of denaturation at 95°C for 15 s and annealing at 55°C for 30 s and extension at 72°C for 30 s. Reactions were performed in triplicate and the mRNA expression was normalized against the endogenous reference (18S). Data were quantified by using CFX96TM real-time PCR detection system (Bio-Rad; Hercules, CA, USA).

### Western Blot Analysis

Tissues or dissociated neurons were collected and lysed using a RIPA buffer for 30 minutes on ice. The protein concentration was measured by using the BCA Kit used according to the manufacturer’s instructions (Beyotime, China). Then, equal amount of extracted proteins were separated with 10% SDS-PAGE, followed by transferring to the polyvinylidene difluoride (PVDF) membrane, Immobilon-P (Millipore; Burlington, MA, USA). Then, PVDF membranes were blocked with 5% non-fat milk in Tris-Buffered Saline (TBS) buffer for 1 hour at room temperature, and reacted with primary antibody overnight at 4°C. Then, the membrane was incubated with horseradish peroxidase-conjugated secondary antibody (dilution of 1:1000) at room temperature for 1 hour. Lastly, the membrane was developed using an ECL Prime Western Blotting Detection Reagent (GE Healthcare; Chicago, IL, USA). The density of protein bands from three independent experiments was quantified using the Image J software (National Institutes of Health [NIH]; Bethesda, MD, USA).

### *in vivo* Electroporation of Adult Mouse DRGs

The *in vivo* electroporation of adult mouse DRGs was performed as described previously (Saijilafu et al., 2012). Briefly, left L4-L5 DRGs of 10-week old adult ICR mice (weighing from 30g to 35g) were surgically exposed under anesthesia. A solution containing siRNAs or c-Myc plasmid and/or EGFP plasmid (1μl) was slowly injected into the DRG using a capillary pipette powered by the Picospritzer III (Parker Inc.; Cleveland, OH, USA; pressure: 30 psi; duration: 8 ms). The electroporation was performed immediately using a tweezer-like electrode (Ø1.2 mm) and an ECM830 Electro Square Porator BTX (five 15 ms pulses at 35 V with 950 ms interval). The mice were then allowed to recover after surgery after the incision sites were closed. The ipsilateral left sciatic nerve was then crushed with fine forceps two days later, and the crush site was marked with an 11-0 nylon epineural suture. Three days after the nerve crush, the mice were terminally anesthetized and transcardially perfused with ice-cold 4% PFA. The entire sciatic nerve was dissected out and further fixed in 4% PFA overnight at 4°C. The epineural suture was confirmed, and the lengths of all EGFP-labeled regenerating axons were measured from the crush site to the distal axon tips. The number of the independent experiments performed and the number of neurons analyzed in each condition are depicted in the Figure or in the legends.

### Sciatic Nerve Axotomy

The sciatic nerve axotomy model was performed as described above. Under anesthesia with ketamine / xylazine, and a 1 cm-long skin incision was made aseptically on the legs. The overlaying muscles were then dissected free from the L4-L5 spinous processes on the two sides. The sciatic nerve was crushed at the sciatic notch with ophthalmic scissors. After a specific period of time, the L4-L5 DRGs were isolated for cell culture, tissue section, qPCR, or western blotting. Mice who received sham surgery were used as “no-injury” controls.

### Intravitreal Injection and Optic Nerve Injury

After mice were anaesthetized, a glass micropipette was inserted into the vitreous body of 4-week old C57 mice avoiding damaging the lens, and a 1 μl of AAV2-GFP or AAV2-p53 (Titer 1×10^12^) viral vector suspension was carefully injected with a Picospritzer III (Parker Inc. pressure: 9 psi; duration: 5 ms). Two weeks after the virus injection, the optic nerve was exposed intraorbitally by blunt dissection, and a crush injury was performed approximately 1 mm behind the optic disc with fine forceps for 2 to 3 seconds.

### RGC Axon Anterograde Labeling and Counting of Regenerating Axons

In order to label RGC axons, twelve days after the optic nerve crush injury, 2 μl of fluorescence Alexa-594 conjugated cholera toxin β subunit (CTB) (2 μg/μl, Invitrogen; Carlsbad, CA, USA) was micro-injected into the vitreous body using a Picospritzer III (Parker Inc.; pressure: 9 psi; duration: 5 ms). Two days later, animals were perfused with 4% PFA in 0.01 M phosphate buffered saline via the left ventricle. The entire optic nerve was then harvested. Regenerating RGC axons were calculated as previously described (Kurimoto et al., 2010). 12-μm thick longitudinal sections from the optic nerves were cut using a cryostat. The numbers of CTB-positive axons were counted in each of the five sections (every four sections) at a different distance from the crush site. The cross-sectional width of the nerve was measured at the counting point and used to calculate the number of axons per mm of nerve width. Then, the average of the number of axons per millimeter on all parts was measured. The total number of axons extending from the nerve of radius “r” was estimated by summing all the parts with thickness t (12 μm): Σad = πr^2^ x [average axons/mm]/t.

### Whole Mount Retina Staining

After perfusion with PBS and ice cold 4% PFA, whole retinas were dissected out and blocked in PBS containing 10% FBS and 0.3% Triton X-100 for 1 hour. Then, incubated with primary antibody Tuj1 (1:500) overnight at 4°C. After washing the retinas with blocking buffer three times for 30 minutes, the tissues were incubated with Alexa Fluor594-conjugated secondary antibodies for 1 hour at room temperature. Whole-mount retina sections were immunostained with a Tuj1 antibody to label RGC neurons. Using Image J software in eight fields per case distributed in four quadrants of the eye, cell survival was reported as the number of Tuj1 positive cells per mm^2^ averaged over the eight fields sampled in each retina and then averaged across all cases within each experimental group. The quantitation obtained from the regeneration and cell survival experiments were based on 6 mice per condition.

### Statistics

All statistical analyses were expressed as the mean ± standard error mean (SEM). To determine significance between two groups, comparisons between means were made with the Student’s *t*-test. Multiple group comparisons were performed with one-way ANOVA analysis of variance using the statistical software Prime 5 followed by Bonferroni’s post-test as post hoc test for comparing multiple groups. Statistical significance was considered when the *p*-value was less than 0.05 for three independent experiments (\**p* < 0.05; \*\**p* < 0.01; \*\*\**p* < 0.001).

## Author Contributions

JJ Ma, RJ Xu, X Ju, WH Wang, ZP Luo, CM Liu, L Yang, B Li, JQ Chen, B Meng, Hl Yang, FQ Zhou, and Saijilafu designed the experiment. JJ Ma, RJ Xu, X Ju and WH Wang performed the experiments and analyzed the data. JJ Ma, FQ Zhou, and Saijilafu co-wrote the paper with all authors’ input.

## Acknowledgments

Dr. Saijilafu was supported by the grant from the National Natural Science Foundation of China (Nos. 81772353 and 81571189), the National Key Research and Development Program (Nos. 2016YFC 1100203), The Priority Academic Program Development of Jiangsu Higher Education Institutions, and Innovation and Entrepreneurship Program of Jiangsu Province. Dr. Feng-Quan Zhou was supported by NIH (R01NS064288, R01NS085176, R01GM111514, R01EY027347), the Craig H. Neilsen Foundation, and the BrightFocus Foundation.

## Supplementary figure legends

**Supplementary Figure S1.**
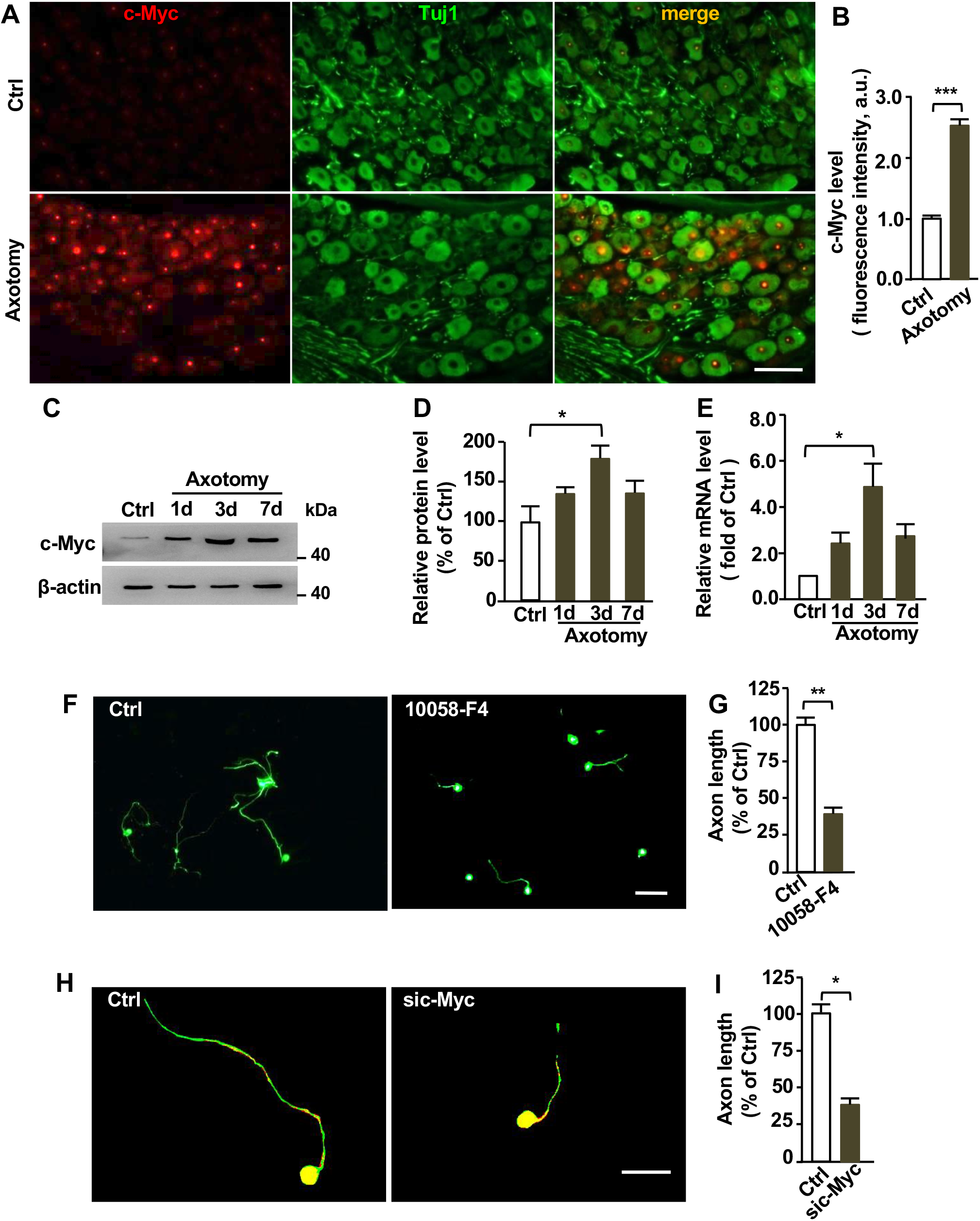
C-Myc is up regulated in adult sensory neurons upon peripheral nerve injury and functionally required for sensory axon regeneration in vitro. (A) Representative immunostaining of DRG sections showing nucleus localization of c-Myc in DRG neurons and up regulation of c-Myc levels after sciatic nerve axotomy. Scale bar: 50 μm. (B) Quantification of nucleus fluorescence intensity of c-Myc (n = 3 independent experiments, \*\*\**p* < 0.001). (C) Representative western blot images showing significantly increased levels of c-Myc in adult DRG neurons *in vivo* 3 days after sciatic nerve axotomy. (D) Quantification of western blot results from three independent experiments; \**p* < 0.05. (E) qPCR results showing significantly increased mRNA levels of c-Myc in adult DRG neurons *in vivo* 3 days after sciatic nerve axotomy, n = 3 independent experiments; \**p* < 0.05. (F) Representative images showing that inhibition of c-Myc in cultured DRG neurons with the pharmacological agent, 10058-F4, for 3 days dramatically reduced DRG axon growth. Scale bar: 200 μm. (G) Quantification of average axon lengths of control neurons and neurons treated with the c-Myc inhibitor from three independent experiments; \*\**p* < 0.01. (H) Representative images showing that knocking down c-Myc with a specific siRNA in cultured DRG neurons dramatically reduced axon growth after 3 days in culture. Scale bar: 50 μm. (I) Quantification of average axon lengths of control neurons and neurons expressing sic-Myc from three independent experiments; \**p* < 0.05.

**Supplementary Figure S2.**
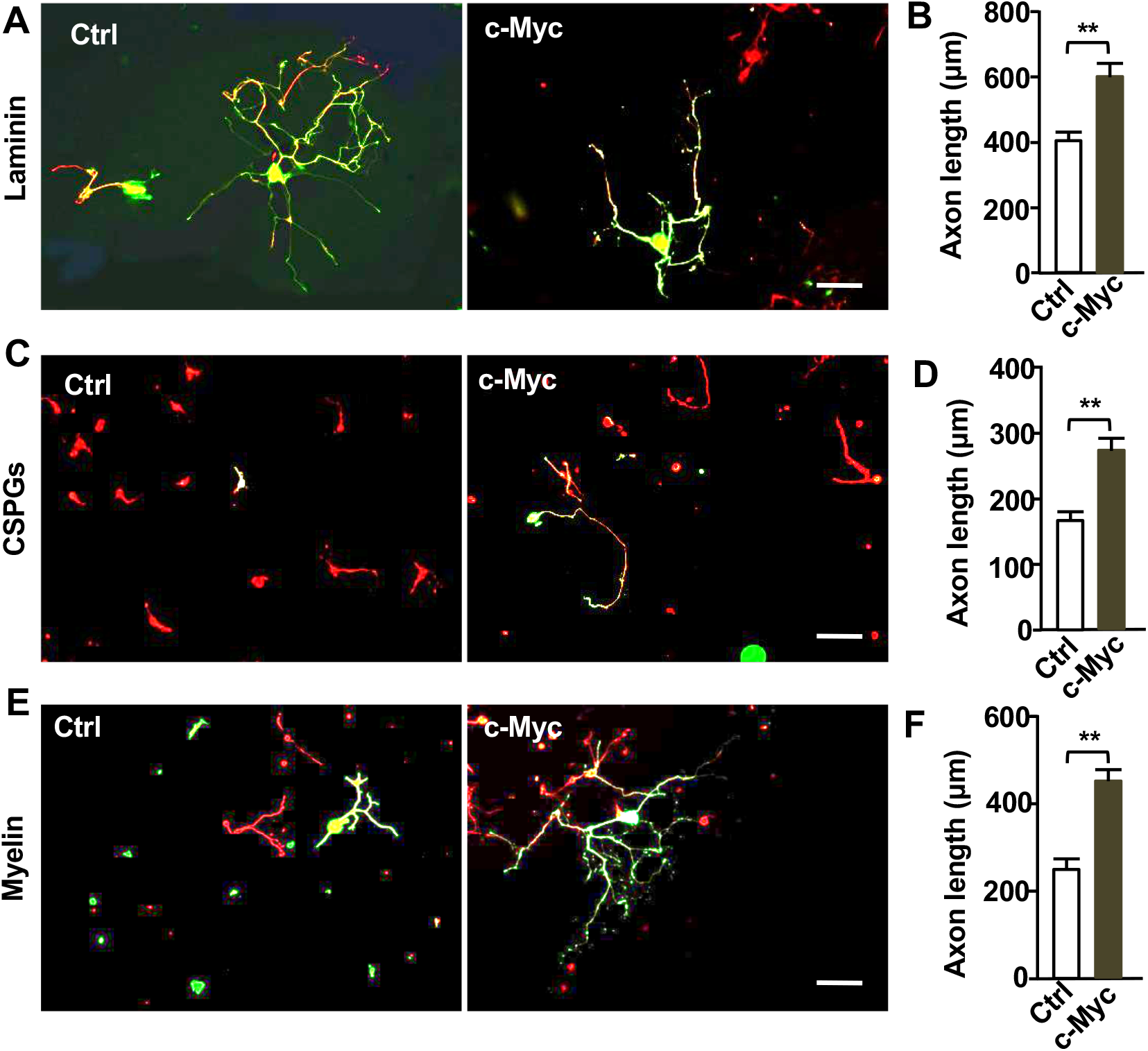
Overexpression of c-Myc in sensory neurons is sufficient to promote axon growth over both supportive and inhibitory substrates. (A) Representative images showing that overexpression of c-Myc significantly enhanced regenerative axon growth of sensory neurons cultured on laminin. Scale bar: 200 μm. (B) Quantification of (A) from three independent experiments; \*\**p* < 0.01. (C) Representative images showing that overexpression of c-Myc significantly enhanced regenerative axon growth of sensory neurons cultured on the inhibitory substrates, chondroitin sulfate proteoglycan (CSPGs). Scale bar: 200 μm. (D) Quantification of (C) from three independent experiments; \*\**p* < 0.01. (E) Representative images showing that overexpression of c-Myc significantly enhanced regenerative axon growth of sensory neurons cultured on myelin inhibitory substrates. Scale bar: 200 μm. (F) Quantification of (E) from three independent experiments; \*\**p* < 0.01.

**Supplementary Figure S3.**
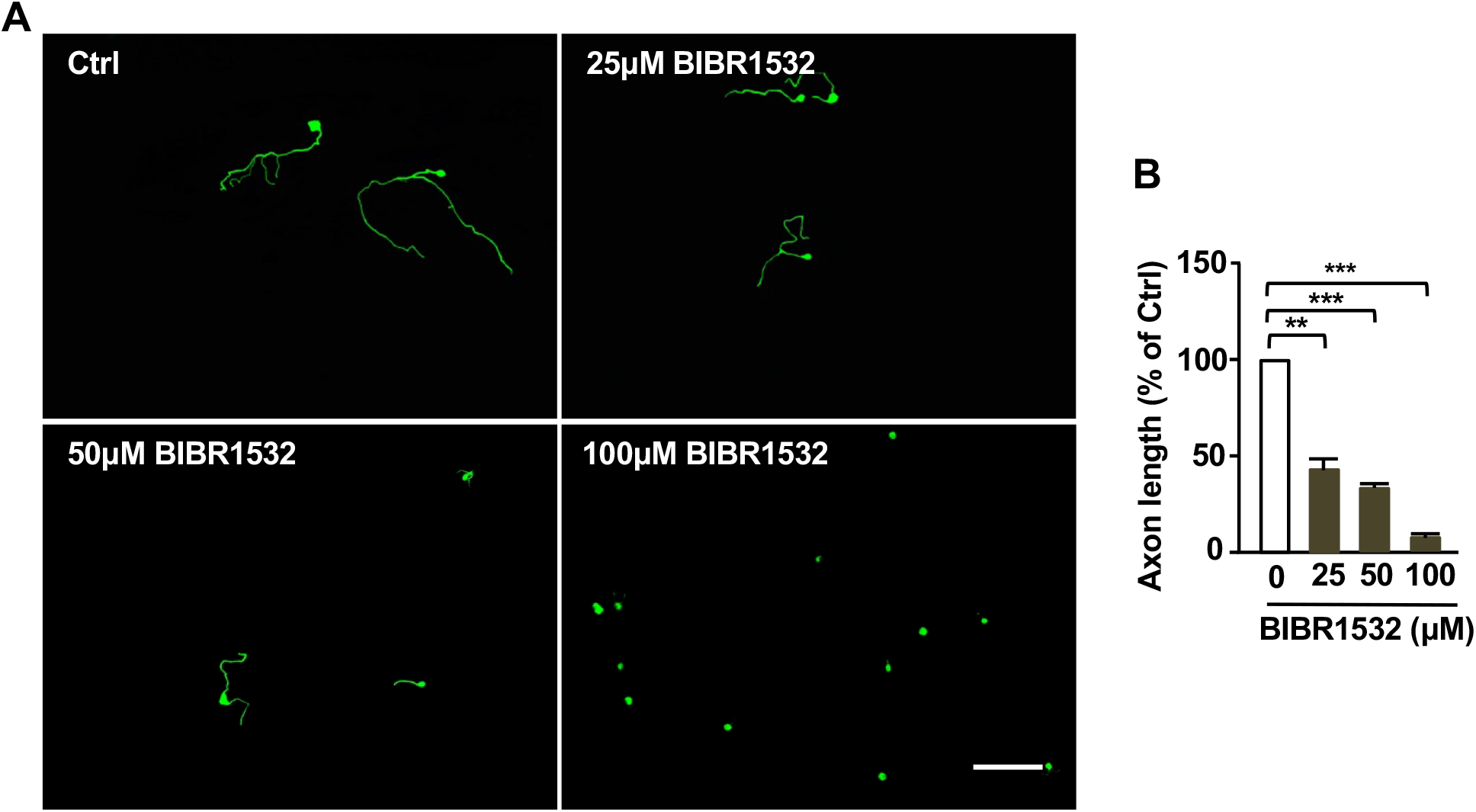
Inhibition of TERT with specific inhibitor BIBR1532 impairs regenerative axon growth of sensory neurons in a dose dependent manner. (A) Representative images showing that inhibition of TERT activity with the pharmacological inhibitor, BIBR1532, at different concentrations, blocked regenerative axon growth of sensory neurons. Scale bar: 200 μm. (B) Quantification of the average axon lengths of control neurons and neurons treated with 3 different concentration of BIBR1532 from three independent experiments; \*\**p* < 0.01, ***p < 0.001.

**Supplementary Figure S4.**
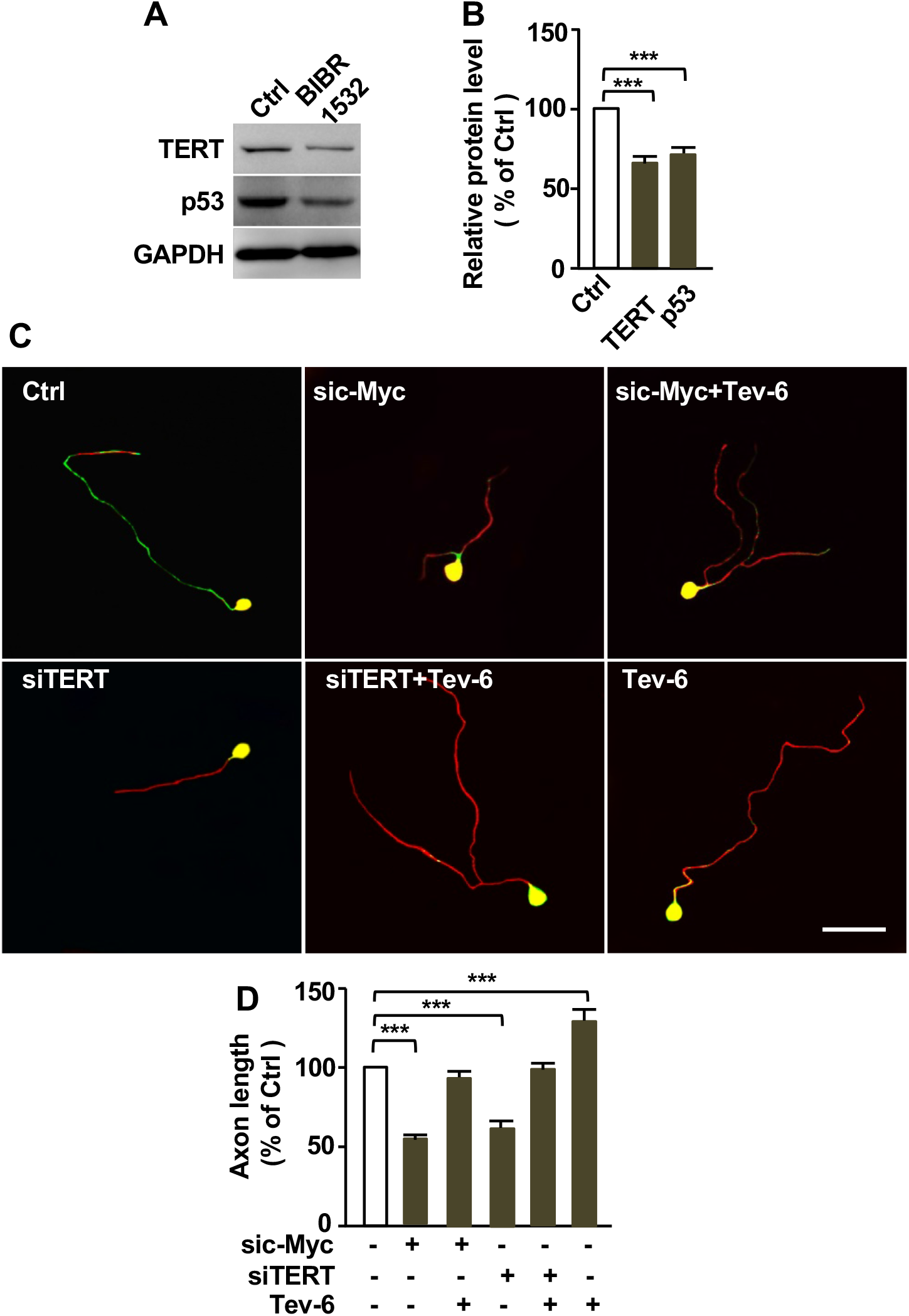
p53 expression is increased in adult mouse sensory neurons upon peripheral nerve injury and its activity is required for regenerative sensory axon growth. (A) Representative western blot images showing that inhibition of TERT activity with its specific inhibitor BIBR1532 led to reduced levels of TERT and p53 in sensory neurons. (B) Quantification of the Western blot images in (A) from three independent experiments; \*\*\**p* < 0.001. (C) Representative images showing that activation of p53 activity with Tenovin-6 (0.5 μM) restored sensory axon regeneration defects induced by knocking down c-Myc or TERT. Scale bar: 50 μm. (D) Quantification of the average axon lengths in (C) from three independent experiments; \*\*\**p* < 0.001.

**Supplementary Figure S5.**
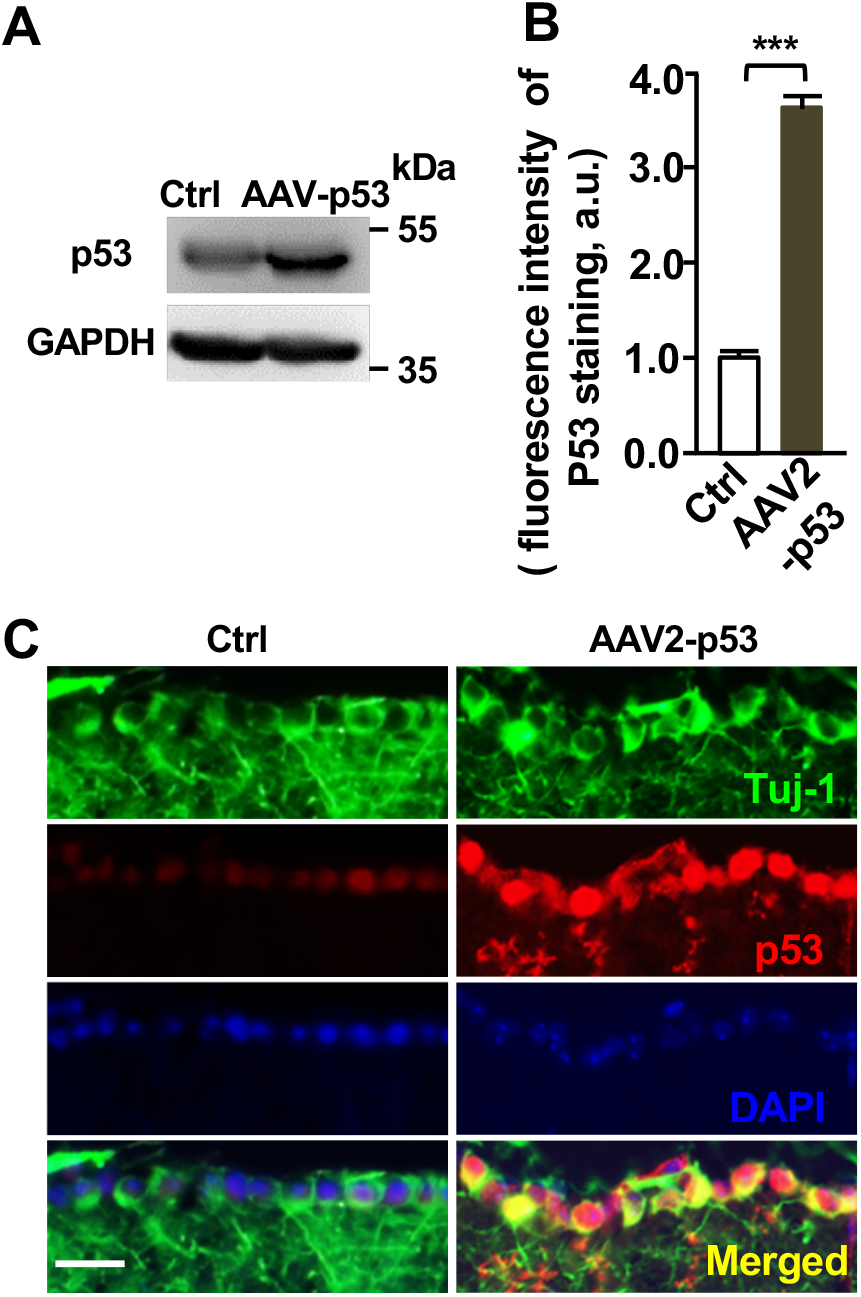
AAV2-p53 mediated overexpression of p53 in retinal ganglion cells (RGCs) (A) Representative western blot images showing increased expression of the p53 protein in mouse retina tissues. (B) Quantification of fluorescence intensity of p53 in RGCs from three independent experiments; \*\*\**p* < 0.001. (C) Representative images of immunostaining of sectioned retina with neuronal marker Tuj-1, p53, and nuclear DNA dye DAPI. Note the markedly increased p53 staining in RGCs 2 weeks after vitreous injection of the adeno-associated virus vector, AAV2-p53. Scale bar: 50 μm.

